# Discovery of a Novel Double-Stranded DNA Virus Reveals its Long-term Co-evolution with Ants

**DOI:** 10.1101/2025.02.14.638241

**Authors:** Shengqiang Jiang, Liangliang Zhang, Xingyu Guo, Jianchao Li, Jing Hu, Hong He, Hongying Chen

**Author notes:** Correspondence: Hongying Chen, (H.C.); Hong He, (H.H.). These authors contributed equally to the work (Shengqiang Jiang, Liangliang Zhang).

## Abstract

Large double-stranded DNA (dsDNA) viruses have been shown to have a wide host range in insects. However, their infection in ants has not yet been described. In this study, a novel filamentous virus, Camponotus japonicus labial gland disease virus (CjLGDV), was identified in the enlarged labial gland of *Camponotus japonicus*. By transmission electron microscopy, the nucleus of infected labial gland cell was shown to be full of non-enveloped nucleocapsids, and enveloped virus particles were observed in the cytoplasm. Genome sequencing analysis shows that CjLGDV possesses a circular dsDNA genome of 142 kb. By sequence alignment, a CjLGDV-like viral genome was found in *Anoplolepis gracilipes*, and conservation of gene synteny was observed between CjLGDV and the CjLGDV-like virus. Phylogenetic analysis suggests that CjLGDV and CjLGDV-like virus represent two members of a novel virus family, distantly related to unclassified Apis mellifera filamentous virus and Apis mellifera filamentous-like virus. These viruses have common genomic characteristics and key conserved genes with the viral members in the order *Lefavirales*, class *Naldaviricetes*. By mining public databases, CjLGDV-related endogenous viral elements (EVEs) were detected in the genome of multiple ant species, and phylogenetic analysis indicated a long-term coevolutionary relationship between the EVEs and its host. These findings expand the known lineage of nuclear arthropod large DNA viruses (NALDVs), broaden our understanding of their host range, and reveal a long and frequent interaction history between CjLGDV and its ant hosts.

## Introduction

Recent advancements in molecular biology and high-throughput sequencing technologies have substantially enhanced our understanding of virus-host interactions and expanded our knowledge of the global virome (Rascovan et al. 2016; Shi et al. 2016; Zhang et al. 2018; Briese et al. 2015). To date, gene sequences of thousands species of viruses have been identified<u> in</u> dozens of ant species (Li et al. 2024; Flynn and Moreau 2024; Baty et al. 2020). However, the infectivity and transmissibility of these viruses in ants remain largely speculative and unconfirmed. Among those confirmed, the majority are single-stranded RNA (ssRNA) viruses belonging to the order *Picornavirales* (Baty et al. 2020). Some of these viruses, such as Solenopsis invicta virus 3 (SINV- 3) which alters foraging behavior and increases brood mortality in its host, have been explored as potential biological control agents targeting the invasive red imported fire ant (Valles et al. 2022; Valles 2023, 2024). Despite extensive research on RNA viruses, documentations of DNA viruses that are infectious for ant are rare. To our best knowledge, the only DNA virus characterized in ant is Solenopsis invicta densovirus (SIDNV), a single-stranded DNA virus found in South American populations of *Solenopsis Invicta* (Valles et al. 2013). Even so, the impact of SIDNV on its host remains unclear. Recent studies on ant viral diversity have uncovered numerous DNA viral genome fragments in several ant species, indicating a broader and more complex landscape of DNA viral infections in ants than previously thought (Flynn and Moreau 2024).

Endogenous viral elements (EVEs), or viral fossils, are whole or fragmented viral sequences integrated into host genomes after the virus infection and preserved in the host through germline transmission. Recent findings on the distribution of EVEs in ant genomes have significantly advanced our understanding of the viral community composition in ant (Flynn and Moreau 2019; Gilbert and Belliardo 2022; Guinet et al. 2023). Notably, EVEs related to large double-stranded DNA (dsDNA) viruses have been identified in several ant species, suggesting a historical prevalence of DNA virus infections.

According to the International Committee on Taxonomy of Viruses (ICTV) report, classified nuclear arthropod large DNA viruses (NALDVs) are categorized into four families: *Baculoviridae*, *Nudiviridae*, *Hytrosaviridae*, and *Nimaviridae* (Walker et al. 2021; Van Oers et al. 2023). Unclassified filamentous viruses infecting *Hymenoptera* insects, such as Apis mellifera filamentous virus (AmFV), Apis mellifera filamentous-like virus (AmFLV) and Leptopilina boulardi filamentous virus (LbFV), share genetic characteristics with these four virus families. However, phylogenetic analyses suggest that these unclassified filamentous viruses belong to new viral families (Kadlečková et al. 2024; Gauthier et al. 2015; Lepetit et al. 2016).

A condition referred to as “labial gland disease”, characterized by swollen labial glands and malformed mesosomas, has been reported in more than ten ant species from Europe, the USA, Japan and China (Elton 1991; Laciny 2021; Zhang et al. 2023). Although the causative agent remains unknown, it has been postulated that the disease may be transmitted by trophallaxis behavior, cannibalism and probably also vertically from queen to offspring (Elton 1991; Laciny 2021). Assumption of a viral pathogen has been proposed to cause the distinct morphological changes, but it has not been confirmed (Elton 1991; Laciny 2021).

Recently, we reported the observation of labial gland enlargement symptom in *Camponotus japonicus*, an ant species widely distributed across East Asia (Zhang et al. 2023). In this study, we identified a novel virus with circular dsDNA genome from the enlarged labial gland, and named it Camponotus japonicus labial gland disease virus (CjLGDV). Morphological characteristics of the virus particles were recorded by transmission electron microscopy. By searching public database, a CjLGDV-like viral genome was identified in *Anoplolepis gracilipes.* Phylogenetic analysis elucidated the relationships between CjLGDV, CjLGDV-like and other known viruses. Furthermore, CjLGDV-related endogenous viral elements were characterized in the genomes of various ant species, providing evidence for the long and frequent interaction history between CjLGDV and its ant hosts. These findings expand our understanding of the host range and evolutionary dynamics of large dsDNA viruses, and provide evidence for the long-standing coevolutionary relationship between CjLGDV and its ant hosts.

## Results

### Observation of virus particles in enlarged labial gland of *Camponotus japonicus*

In our recent report, we found that about one fifth of the minor workers in a mature nest of *C. japonicus* had swollen labial glands (Fig. 1A) (Zhang et al. 2023). To investigate the cell structure in the enlarged labial gland, the tissue was sliced and examined using transmission electron microscopy (TEM). Numerous long filaments, likely viral nucleocapsids in processing, were observed in the nuclei of the labial gland cells (Fig. 1B). In the cytoplasm, some nucleocapsids appeared coiled within spherical enveloped vesicles around 200 nm, while others were in enveloped filamentous forms measuring up to 1000 nm in length and about 65 nm in width (Fig. 1C-D). Transitional stages between these two morphologies were also observed. These observations suggest that the causative agent of the swollen labial gland is probably an enveloped filamentous virus replicated in the nucleus, which is named as Camponotus japonicus labial gland disease virus (CjLGDV).

**Fig. 1.**
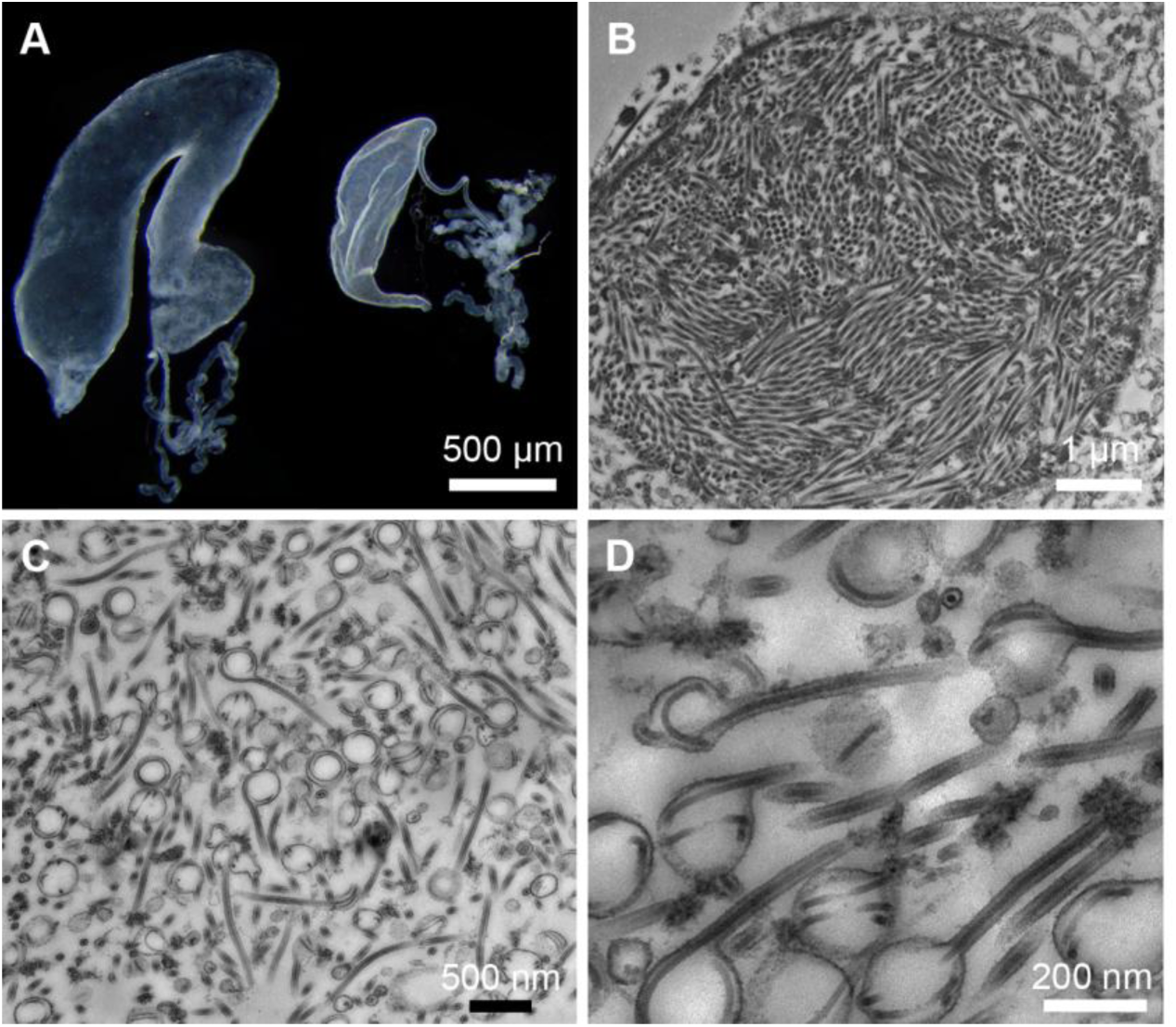
Transmission electron micrographs of Camponotus japonicus labial gland virus (CjLGDV) in enlarged labial gland of *C. japonicus*. (A) Normal (right) and enlarged labial glands (left) of *C. japonicus.* (B) The cell nucleus in an enlarged labial gland of *C. japonicus* is full of nucleocapsid-like filamentous particles. (C) Enveloped nucleocapsids in the cytoplasm. (D) A high magnification image of the enveloped CjLGDV in the cytoplasm

### General features of the viral genome

Based on the observation of numerous nucleocapsid-like filaments in the cell nucleus and the fact that most DNA viruses replicate in the nucleus, we speculated that the causative agent of the labial gland disease could be a DNA virus. To confirm this speculation, DNA was respectively extracted from the normal labial gland (LG) and enlarged labial gland (ENLG) of *Camponotus japonicus* and subjected to Illumina sequencing. For the LG group, 91.7% of the reads were mapped to the *Camponotus* genome, while only 48.9% reads were mapped to the ant genome in the ENLG group (Supplementary Fig. S1).

The unmapped ENLG reads were used for viral genome assembly. After removing low-abundance contigs and assembly of overlapping ones, ambiguous and gap regions were resolved by PCR amplification (Supplementary Fig. S2) and Sanger sequencing. The primers for the PCR reactions, and the length and position of the fragments are listed in Supplementary Table S1. Finally, we obtained a 142,484 bp circular dsDNA genome (Fig. 2A), which falls within the size range of arthropod large dsDNA viruses. General features of GjLGDV genome in comparison with other arthropod large dsDNA viruses are listed in Table 1. The GC content in GjLGDV genome is moderate at 52%. In contrast, most baculoviruses, nudiviruses, and hytrosaviruses with similar genome sizes have much lower GC contents (Table 1).

**Fig. 2.**
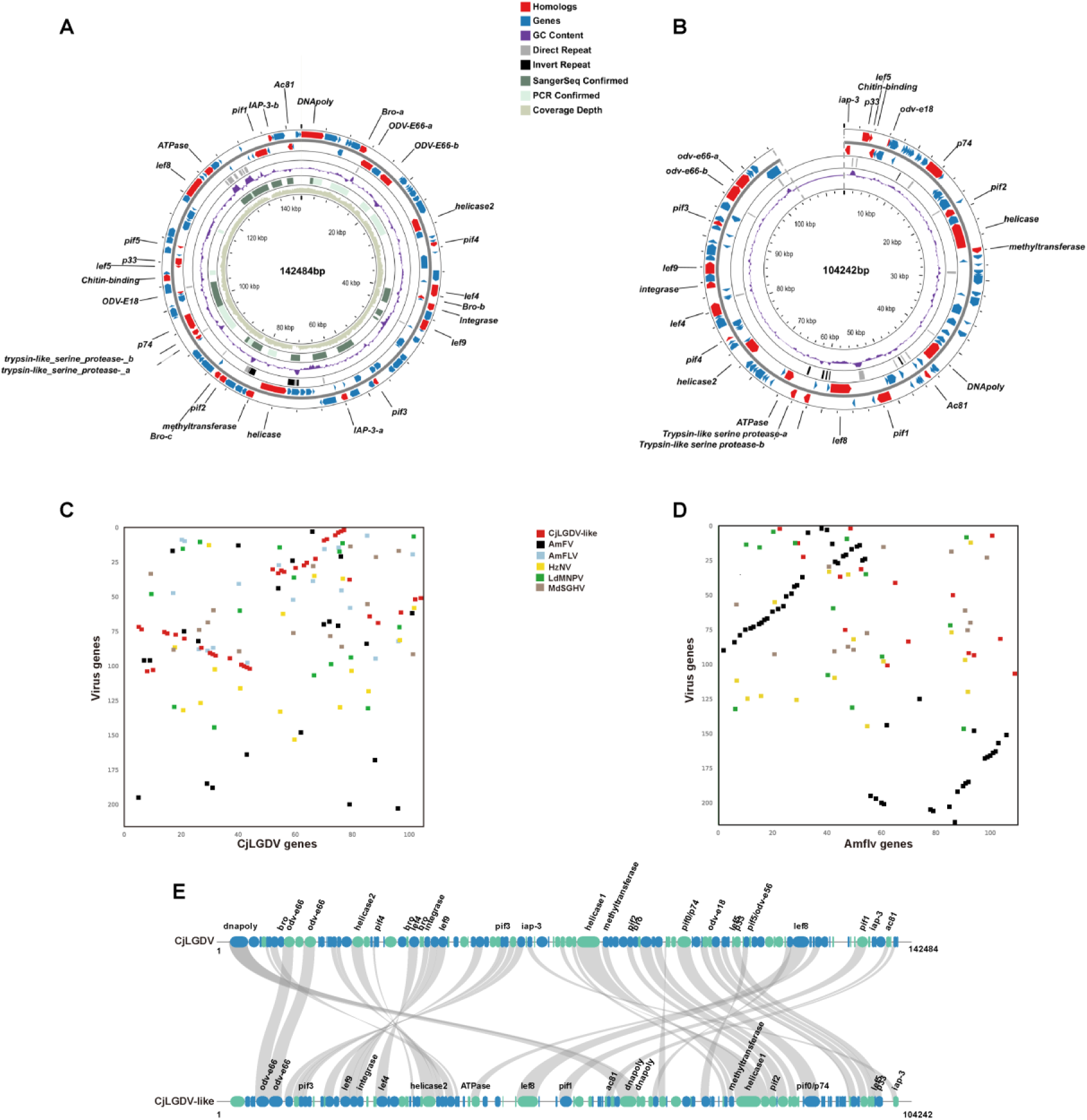
Characterization of CjLGDV and CjLGDV-like genomes. Diagrams of the CjLGDV (A) and CjLGDV-like (B) genome. Putative ORFs and their directions are indicated by arrows. Red arrows represent the ORFs having homologous genes within *Naldaviricete*, and the gene names are labeled in the diagram. Black rectangles represent inverted repeats, grey rectangles represent direct repeats, green rectangles represent PCR confirmed regions, and dark green rectangles represent regions confirmed by Sanger sequencing. The GC content along the genome is displayed in purple. The inner gray green circle indicates the sequencing coverage depth of the CjLDGV genome. The adenine of the starting codon for the DNA polymerase is set as starting site for the CjLGDV circular genome. (C-D) Gene-order conservation between viruses. Each gene is represented by a dot, with the x-axis showing its order in the CjLGDV (C) or AmFLV (D) genome and the y-axis indicating the position of its homolog in other virus genomes. (E) Gene synteny between CjLGDV and CjLGDV-like. Homologs between CjLGDV and CjLGDV-like are linked by gray lines. Blue and green ellipses represent genes on the positive and negative strand, respectively.

**Table 1.**
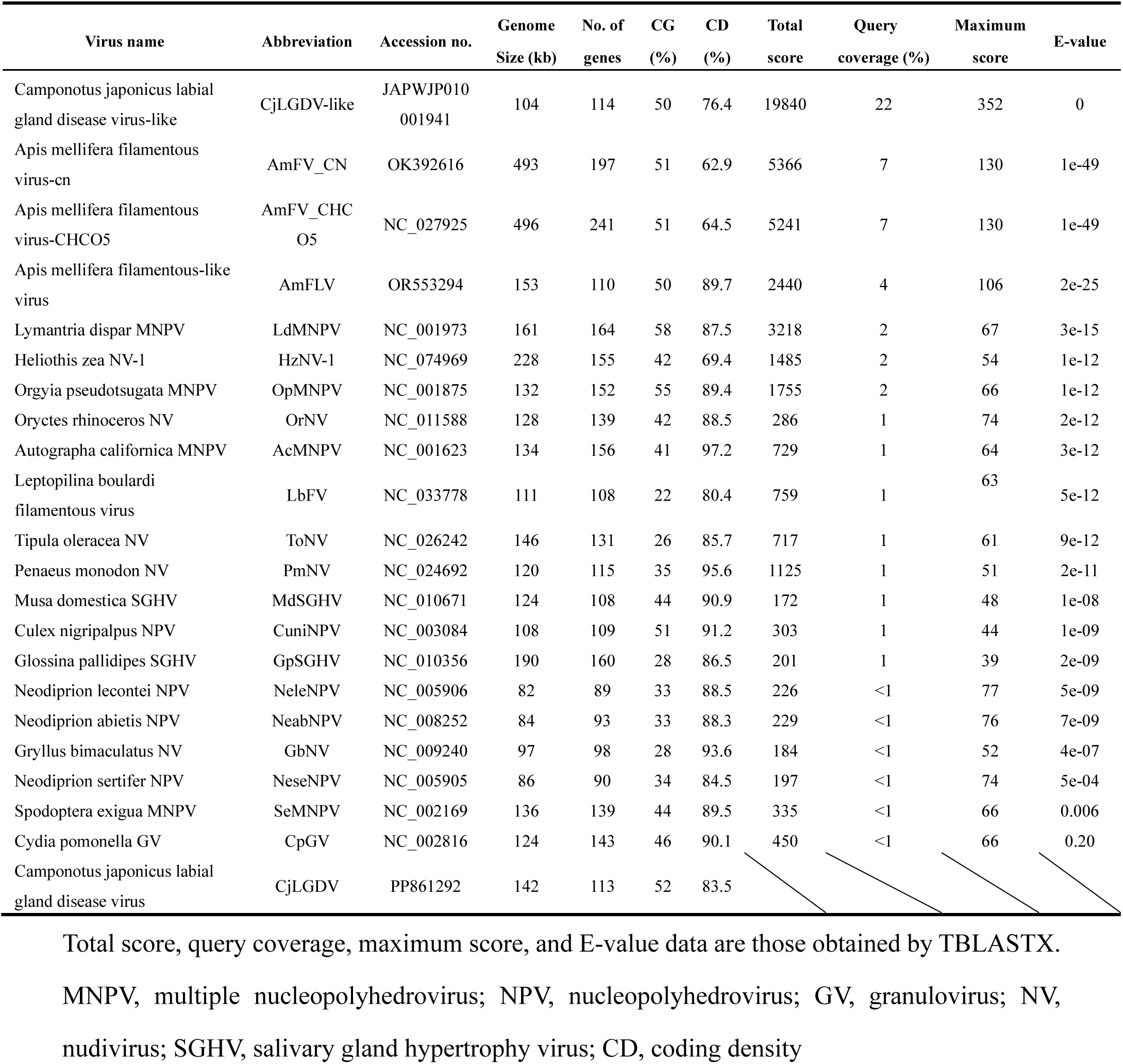
General features of GjLGDV genome in comparison with other arthropod large dsDNA viruses.

Given the possibility that CjLGDV may infect multiple ant species, or that related viruses may exist, genetic sequences similar to CjLGDV could be present in ant sequencing data as a result of viral infections. Genome assemblies of 59 ant species from 6 subfamilies were found in public databases and screened for similar viral sequences by BLAST. Exogenous CjLGDV associated scaffolds were expected to fall within the genome size range of large arthropod dsDNA viruses and exhibit distinct sequence characteristics with host genome scaffolds. Finally, a 104,242 bp scaffold (JAPWJP010001941.1) in the genome assembly of *Anoplolepis gracilipes* (GCA_031304115.1) was identified as a CjLGDV-like sequence (Fig. 2B). Due to the unavailability of the original sequencing data of the assembly, the sequencing depth of the virus-like scaffold was assessed in other reported *Anoplolepis gracilipes* samples (SRR21231523, SRR21232721, SRR21232722), in which no such sequence was discovered. The absence of the scaffold in other sequencing samples suggests that it only originates from infected samples. The scaffold lacks eukaryotic genes and exhibits a distinct GC content compared to host BUSCO scaffolds, further supporting its exogenous origin (Supplementary Fig. S3).

A TBLASTX similarity assessment revealed that CjLGDV shares more similarity with the CjLGDV-like scaffold (22% coverage) and two honey bee filamentous viruses (AmFV, 7% coverage; AmFLV, 4% coverage) than any other viruses (Table 1). A total of 113 and 114 open reading frames (ORFs) with ATG start codons were predicted in the CjLGDV genome and the CjLGDV-like scaffold, respectively. The average lengths of the predicted ORFs were 351 amino acids (aa) for CjLGDV and 232 aa for the virus-like scaffold, with coding densities of 83.5% and 76.4%, respectively. These genome features are comparable to those of other arthropod large dsDNA viruses with similar genome sizes (Table 1). Among the predicted ORFs, 38 exhibit BLASTP sequence similarities between CjLGDV and the CjLGDV-like scaffold, with amino acid identities ranging from 23.9% to 70.7% (Supplementary Table S2A). Notably, a high level of gene synteny was observed between the two viruses, and the feature was absent when the CjLGDV genome was compared with other viruses (Fig. 2C-E). These findings suggest that the CjLGDV- like scaffold is sufficiently complete to represent the origin virus and is closely related to CjLGDV.

### Repeat regions as a common feature

Homologous regions (*hrs*), containing repeated sequences composed of imperfect palindrome sequences, are a common feature found in invertebrates dsDNA viral genomes. In baculoviruses, *hrs* function as origins of virus DNA replication and transcription enhancers (Kool et al. 1995; Leisy and Rohrmann 1993; Theilmann and Stewart 1992). In the CjLGDV and CjLGDV-like genomes, there are 17 and 14 tandem direct repeat (*dr*) sequences (Supplementary Table S3), accounting for 4.4% and 3.6% of their genome, respectively. Repeats in CjLGDV-like are scattered across the genome, whereas in CjLGDV, six repeats are notably clustered in the region 126181-131264 (Fig. 2A, Supplementary Table S3). All repeats harbor clusters of imperfect palindrome motifs (IPMs) (Supplementary Fig. S4). These repeats are highly conserved within each viral genome, but no similarities were detected between CjLGDV and CjLGDV-like or with any other viruses.

Inverted repeat (*ir*) sequences were detected in both viral genomes (Fig. 2A-B, Supplementary Table S3). The paired repeats exhibit over 98% sequence similarity, with CG content ranging from 46% to 60%. Interestingly, the length of inverted repeats in CjLGV-like range from 131 to 312bp, whereas in CjLGDV, repeats are significantly longer, approximately 2,100 bp. The role of these repeats in virus replication remains to be elucidated. In summary, the presence of repeat sequences appears to be a common characteristic of the two ant dsDNA viruses, and the sequences of the repeats are largely virus-specific.

### CjLGDV and CjLGDV-like encode conserved core genes in *Naldaviricetes*

Functional annotations of ORFs in the CjLGDV and CjLGDV-like genomes were conducted by similarity searches based on amino acid sequences and protein structures. In results, 35 ORFs in CjLGDV (Supplementary Table S2B) and 28 in CjLGDV-like (Supplementary Table S2C) were found to have homologs in other large arthropod dsDNA viruses. The remaining ORFs exhibit either low or no similarity to sequences in available databases. Based on the assumption that viral homologs share similar functions, it was predicted that these genes are involved in DNA replication and processing (*DNA pol*, *helicase*, *helicase2*, *integrase*), transcription and processing (*lef*4, *lef5, lef8, lef9, methyltransferase*), viral packaging and morphogenesis (*ac81*, *p33*, *atpase, trypsin-like serine protease,odv-e18*), viral infectivity (*pif0/p74*, *pif1*, *pif2*, *pif3*, *pif4*, *pif5/odv-e56, chitin-binding protein-like, odv-e66*), and apoptosis inhibition (*iap*) (Supplementary Table S2B- C).

Both CjLGDV and CjLGDV-like genomes contain six core genes shared by *Naldaviricetes*, including genes for per os infectivity factors (*pif0, pif1, pif2, pif3*), DNA polymerase gene (*DNA pol*) and sulfhydryl oxidase gene (*p33*) (Fig. 3). The *pif* gene family is essential for oral infectivity and is recognized as conserved core genes of the nuclear arthropod large dsDNA viruses (Van Oers et al. 2023). Except for *pif4*, which is absent in hytrosaviruses and LbFV, five *pif* (*pif0/p74*, *pif1*, *pif2*, *pif3*, *pif5/odv-e56*) genes are present in the genomes of all sequenced viruses in *Naldaviricetes*. CjLGDV encodes all the six *pif* genes in its genome, whereas CjLGDV-like lacks *pif5*.

**Fig. 3.**
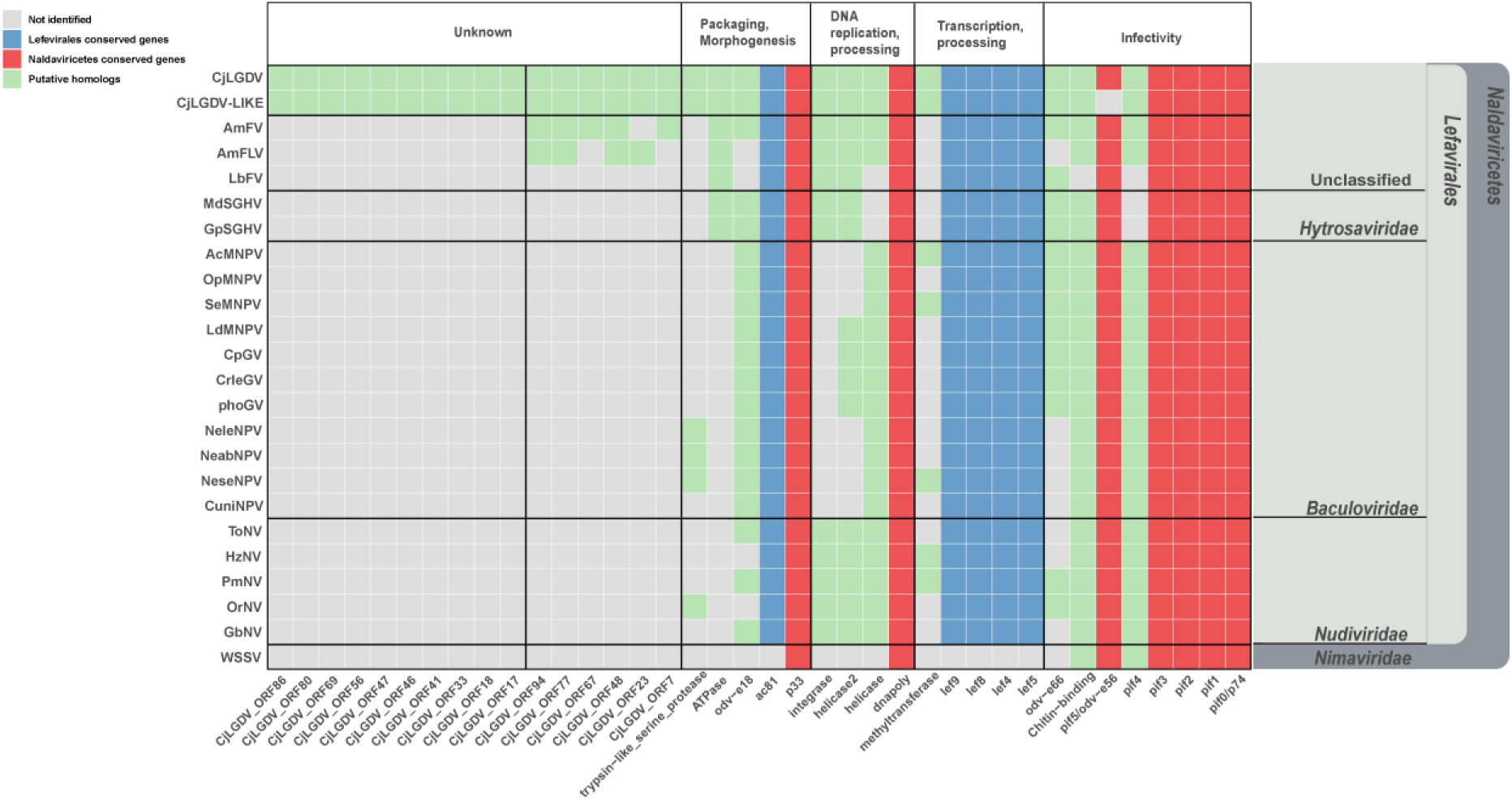
Heatmap of the homologous genes among CjLGDV, CjLGDV-like, unclassified filamentous viruses and representative viruses in *Naldaviricetes.* The taxonomic affiliation of the viruses is marked on the right. Rows represent the viral species, and the columns represent the genes grouped by their potential functions. Colored cells indicate the presence of the gene homolog in viral genomes.

Among the annotated genes, four genes (*DNA pol*, *helicase*, *helicase2*, *integrase*) are identified to be involved in viral DNA replication and processing. Both CjLGDV and CjLGDV-like are predicted to encode a type B DNA polymerase, a common feature of large dsDNA viruses. *Helicase* genes, which are commonly found in members of *Lefavirales*, are also present. Similar to nudiviruses, AmFV, AmFLV and some baculoviruses, two types of *helicase* genes are identified in CjLGDV and CjLGDV-like genomes. The *integrase* is a conserved core gene in nudiviruses and bracoviruses, and it is involved in the excision and circularization of bracovirus DNA (Burke et al. 2013).

*Lefavirales,* comprising the virus families *Baculoviridae*, *Nudiviridae* and *Hytrosaviridae,* is characterized by the possession of conserved baculovirus transcription gene homologs (*lef4*, *lef8*, *lef9*) and can be phylogenetically distinguished from *Nimaviridae* (Van Oers et al. 2023). Five genes (*lef4, lef5, lef8, lef9*, and *ac81*) conserved in *Lefavirales* are detected in both CjLGDV and CjLGDV-like genomes (Fig. 3). In baculovirus, *lef4*, *lef8*, *lef9* and *p47* encode the four subunits of viral RNA polymerase (Guarino et al. 1998; Jin et al. 1998). The homolog of *p47* is absent in CjLGDV and CjLGDV-like, a pattern also observed in hytrosaviruses and honeybee filamentous viruses (Kadlečková et al. 2024; Gauthier et al. 2015; Garcia-Maruniak et al. 2008; Abd-Alla et al. 2008). Additionally, we annotated several previously unidentified *lef* gene homologs in AmFV and AmFLV including *lef4* in AmFV, and *lef4*, *lef5*, *lef8* and *lef9* in AmFLV, based on their sequence similarities to CjLGDV (Fig. 3, Supplementary Table S4).

### Other homologs shared by arthropod large dsDNA viruses

CjLGDV and CjLGDV-like are predicted to encode a FtsJ-like methyltransferase, which is also found in some nudiviruses and baculoviruses (Fig. 3). The methyltransferase is reported to be expressed during the late phase of AcMNPV infection and is involved in the RNA capping process (Wu and Guarino 2003). Its removal, however, does not impact the production of budded or occluded viruses in AcMNPV (Ke et al. 2011). Phylogenetic analysis suggests that viruses may acquire the methyltransferase gene from eukaryotic hosts by horizontal gene transfer (HGT) (Supplementary Fig. S5A).

A putative ATPase from the AAA+ superfamily is detected in CjLGDV and CjLGDV-like, which is also conserved in AmFV, AmFLV, LbFV and hytrosaviruses (Fig. 3). The AAA+ superfamily of ATPases is widely distributed, where its members participate in diverse cellular processes, including membrane fusion, proteolysis and DNA replication (Ogura and Wilkinson 2001; Snider et al. 2008). In baculoviruse infection, host ATPases play critical roles in the construction of the viral replication factory and virion morphogenesis (Li et al. 2020). Phylogenetic analysis suggests that the viral ATPase was more likely acquired somehow from bacteria other than their eucaryotic host (Supplementary Fig. S5B).

Both CjLGDV and CjLGDV-like are predicted to encode a protein containing a chitin-binding domain. Chitin-binding domains are also present in proteins of entomopoxviruses and baculoviruses, which are crucial for the virus oral infectivity (Takemoto et al. 2008; Zhu et al. 2013).

Several multigene families are identified in CjLGDV and CjLGDV-like genomes. Three BRO (Baculovirus Repeated ORFs) homologs with an N-terminal DNA-binding domain are detected in CjLGDV (Supplementary Table S2B). The *bro* genes are prevalent in various arthropod large dsDNA viruses, with baculoviruses containing 0 to 16 *bro* genes. However, the specific functions of these genes still remain unknown.

Insect DNA viruses commonly interfere with the host immune system by preventing apoptosis. Both CjLGDV and CjLGDV-like encode proteins containing the baculoviral IAP repeat (BIR) domain, predicted to be homologs of inhibitors of apoptosis (IAPs). IAPs are anti-apoptotic regulators that prevent apoptosis by inhibiting the caspase family of proteases, and they are widely distributed in large dsDNA virus, yeast, nematodes, insects and mammals (Dubrez-Daloz et al. 2008). Phylogenetic analysis indicates that the ant virus *iap* genes may have eukaryotic origins (Supplementary Fig. S5C).

Two ORFs encoding putative trypsin-like serine proteases are conserved in CjLGDV and CjLGDV-like. Trypsin-like serine proteases are enzymatic proteins commonly found in various RNA and DNA viruses as well as in cellular organisms (Shi et al. 2016). Typically classified as nonstructural proteins in many viruses, these proteases play an indispensable role in viral maturation by facilitating proteolysis through serine-type endopeptidase activity (Bazan and Fletterick 1988; Babéand Craik 1997).

Two baculovirus envelope structural protein homologs (ODV-E18, ODV-E66) are detected in CjLGDV and CjLGDV-like. *odv-e18* is one of the conserved core genes in baculoviruses and essential for the budded viruses production (2008). *odv-e66*, identified in alphabaculovirus and betabaculovirus that infect Lepidoptera hosts, is also present in hytrosaviruses, LbFV, AmFV and certain nudiviruses. The copy number of *odv-e66* varies among viruses, with most containing 0-5 copies. However, in bracoviruses, it has expanded to 36 genes distributed across 10 genomic regions, likely playing a critical role in wasp adaptation (Gauthier et al. 2021). In baculoviruses, ODV-E66 is a major envelope protein with chondroitinase activity that degrades the larval peritrophic membrane, facilitating oral infection (Hou et al. 2019). In the CjLGDV and CjLGDV- like genomes, two adjacent *odv-e66* homologs are annotated (Fig. 2A-B). Phylogenetic analysis indicates that ant virus *odv-e66* homologs cluster with those of NALDVs, forming a monophyletic subclade from bacteria, suggesting that these viral *odv-e66* genes might be acquired by an ancestral virus from bacteria (Supplementary Fig. S6A). Additionally, we found that the viral ODV-E66 homologs share a conserved domain with bacterial chondroitin lyases, including three conserved amino acids previously identified crucial for the enzyme activity in bacteria and baculoviruses (Lin et al. 1997; Rodrigues et al. 2020) (Supplementary Fig. S6A). Of the two ant viral ODV-E66 homologs, ODV-E66a retains all the three conserved amino acids, while ODV- E66b is mutated at two of the three sites (N to D and H to R). The mutations at the conserved sites are predicted to cause some structural changes in the enzyme activity center which may affect the function of the protein (Supplementary Fig. S6B).

### Phylogenetic position of CjLGDV and CjLGDV-like

CjLGDV shares several key characteristics with the viruses in *Naldaviricetes*, including their infection of arthropod hosts, the presence of rod-shaped nucleocapsids in envelope, large circular dsDNA genome and conserved core genes. These shared traits strongly support the classification of CjLGDV as a member of *Naldaviricetes*. To elucidate the phylogenetic relationships of CjLGDV and CjLGDV-like, a highly supported phylogenetic tree was generated using maximum likelihood method, based on the concatenated alignment of the 12 genes conserved in *Lefavirales* (Fig. 4). In this tree, the interrelationships between baculovirus, nudivirus, hytrosavirus and bracovirus families are consistent with previously reported results (Bézier et al. 2014; Kawato et al. 2019). CjLGDV and CjLGDV-like cluster together, occupying a unique position within the phylogenetic tree. They are grouped with AmFV and AmFLV, forming a distinct clade adjacent to LbFV and hytrosaviruses, but far from baculoviruses and nudiviruses. Evolutionary distances within and between virus families in the *Lefavirales* were calculated (Supplementary Fig. S7). As expected, patristic distances within families are smaller (0.12 to 3.06) than those between families (4.21 to 5.01). The distance between CjLGDV and CjLGDV-like viruses (1.19) fall within the range observed for intrafamily distances, being significantly lower than interfamily distances. Interestingly, the distances between honeybee filamentous viruses (AmFV and AmFLV) and ant viruses (CjLGDV and CjLGDV-like) (2.69 to 2.83) are close to the upper limit but still fall within the range of intrafamily distances, suggesting their close evolutionary relationship and these unclassified viruses may belong to a common new virus family.

**Fig. 4.**
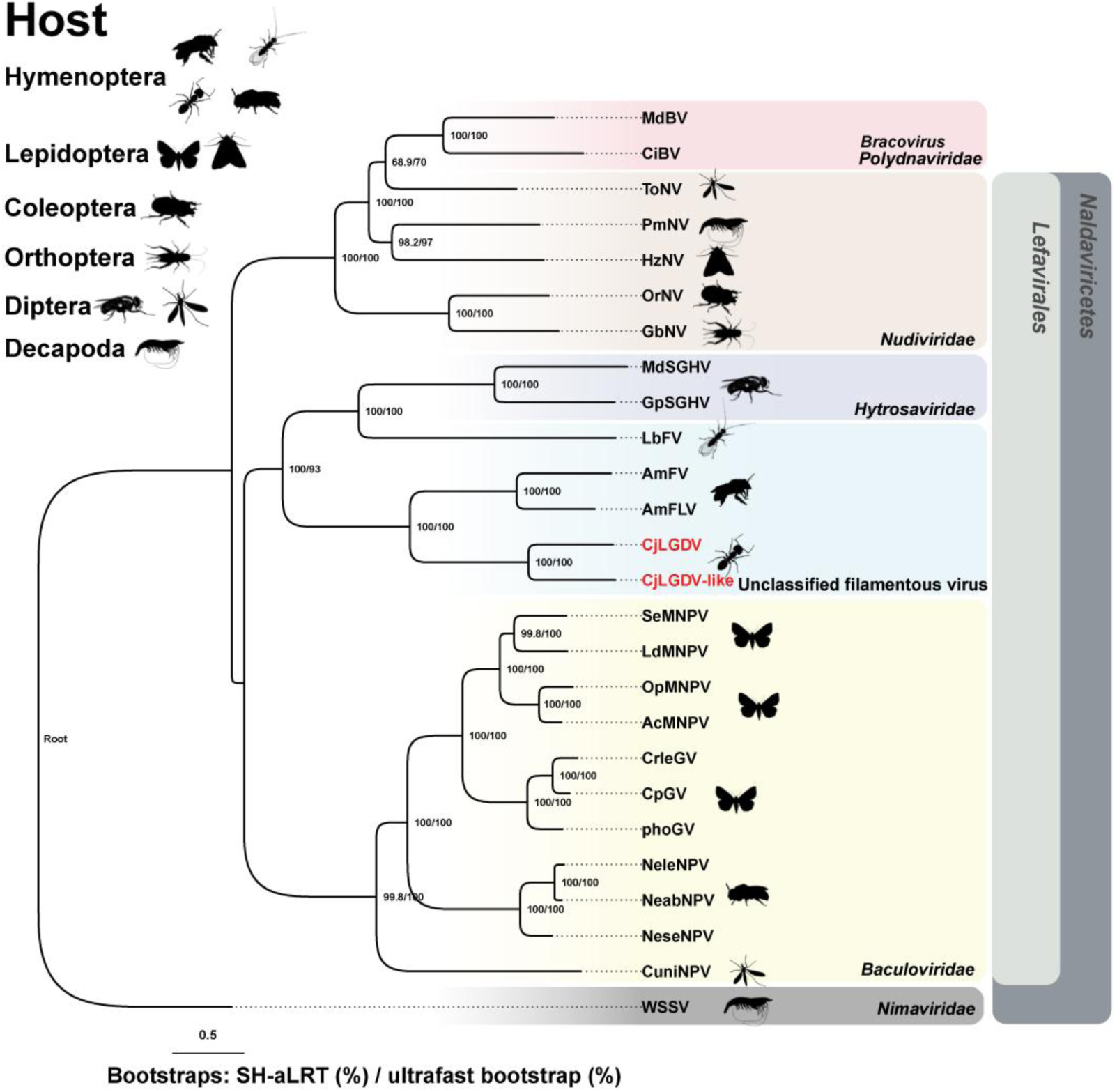
Phylogeny of CjLGDV with arthropod large dsDNA viruses. The phylogenetic tree was constructed using maximum likelihood (ML) inference based on concatenated amino acid sequences of 12 conserved genes (*p74*, *pif1*, *pif2*, *pif3*, *pif5*, *lef4*, *lef5*, *lef8*, *lef9*, *dnapoly*, *p33*, *ac81*). Gene accession numbers are provided in Supplementary Table S4. Node support values are indicated as SH-aLRT support (%) / Ultrafast bootstrap (%). The scale bar represents the average number of amino acid substitutions per site across the tree. Each viral family is denoted by a unique color, and icons next to each virus name indicate the arthropod order of the respective host.

Overall, the phylogenetic analysis reveals a close relationship between CjLGDV and CjLGDV- like, and they may represent members of a novel virus clade in the class *Naldaviricetes,* order *Lefavirales*.

### Endogenous viral elements in ant genomes

Previous researches have demonstrated a rich diversity of endogenous viral elements (EVEs) in ant genomes, highlighting their evolutionary and functional significance (Flynn and Moreau 2019; Guinet et al. 2023). To expand our understanding of ant EVEs, we performed similarity searches against the genomes of 59 ant species available in NCBI databases, using CjLGDV genome as a query.

A total of 1843 loci were identified as candidate EVEs across the genomes of 35 ant species from five subfamilies, including *Myrmicinae*, *Formicinae*, *Ponerinae*, *Dolichoderinae* and *Pseudomyrmicinae* (Supplementary Table S5). Of these, 1705 loci were considered high confidence endogenous sequences, supported by the presence of transposable elements, insect genes, premature stop codons, and similar sequencing depth with scaffolds containing Benchmarking Universal Single-Copy Orthologs (BUSCO) genes (Supplementary Fig. S8, Supplementary Table S5). In a previous comprehensive study of insect EVEs, 278 high-scoring AmFV-related EVEs were identified in ants (Guinet et al. 2023). Among these, 248 were re-identified in our results (Supplementary Table S5), highlighting the sensitivity and accuracy of the detection pipeline.

The EVEs identified in this study exhibit homology to 54 CjLGDV genes, including 21 genes with homologs in NALDVs, 6 genes with homologs in AmFV and AmFLV, 9 genes with homologs in CjLGDV-LIKE, and 18 genes unique to CjLGDV (Fig. 5). BLASTP analysis revealed amino acid sequence identities between the EVEs and viral homologs, ranging from 22% to 76.9% (Supplementary Table S5). Phylogenetic analyses demonstrated a closer relationship between these EVEs and CjLGDV or CjLGDV-like than with AmFV, AmFLV, or other known viruses (Supplementary Fig. S9 1-28). These EVEs are distributed across various scaffolds in ant genomes, with scaffold sizes ranging from hundreds of base pairs to hundreds of millions of base pairs (Supplementary Fig. S10). Each scaffold contains between 1 to 37 viral homologs.

**Fig. 5.**
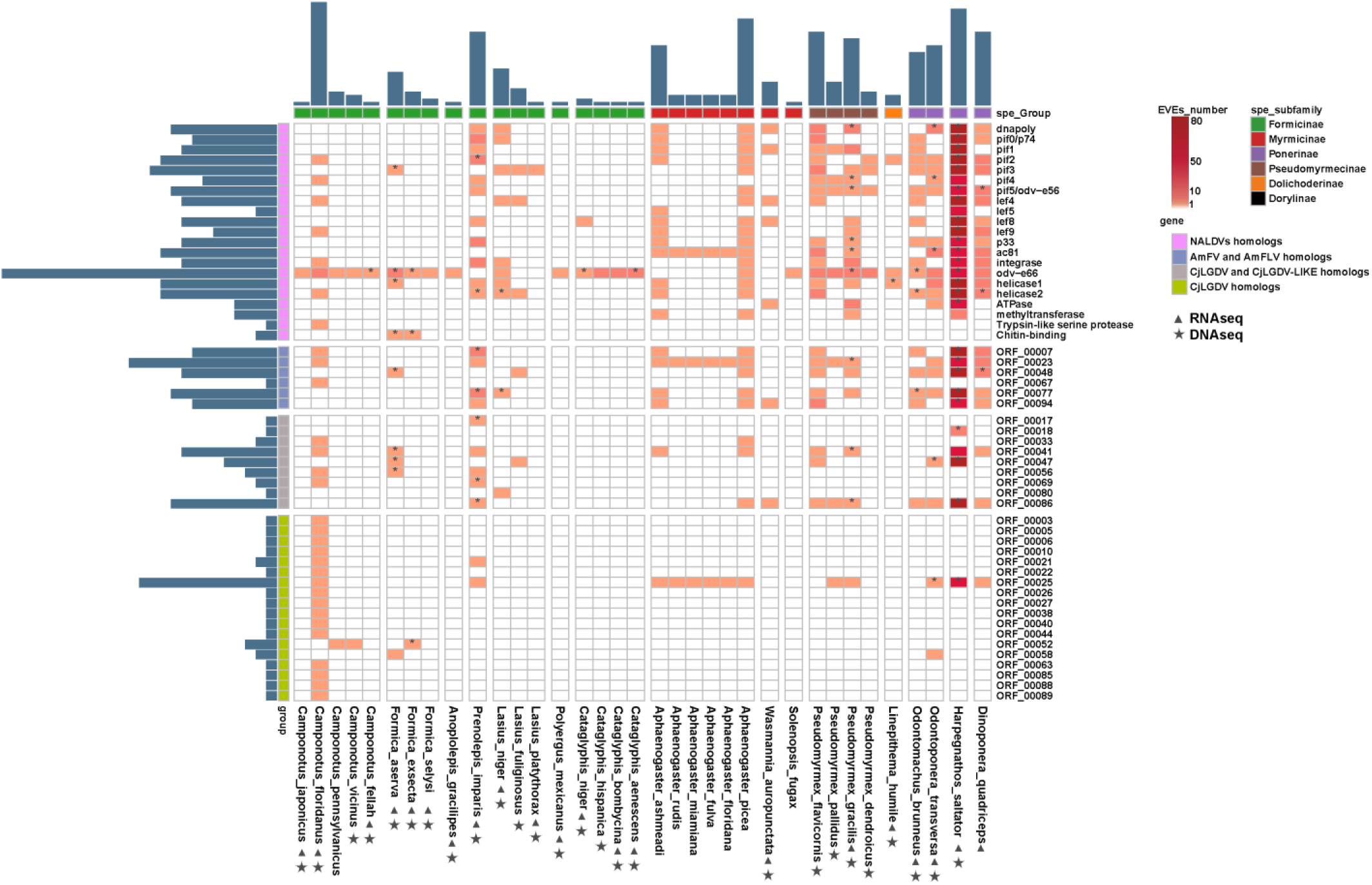
CjLGDV-related endogenous viral elements in ant genomes. Endogenous viral elements identified in the genomes of 35 ant species are shown in a heatmap. Orange cells indicate the presence of the viral homolog in ant genomes. Rows represent the viral homologs, and columns represent the ant species. Bar plots adjacent to the heatmap represent the number of homolog types in each ant (top) and the number of ant species containing each viral homolog (left). Cells marked with * indicate that the gene expression has been detected in the ant. Filled triangle and star under the ant names respectively indicate that raw RNA seq and DNA seq data of that species are available in databases.

Among the ant species analyzed, the highest abundance of EVEs was detected in *Harpegnathos saltator*, where 1378 loci spread across 174 scaffolds showed homology to 29 CjLGDV genes. These homologs present numerous paralogs in the ant genome (Fig. 5), consistent with a previous observation (Guinet et al. 2023). In *Camponotus floridanus*, a distinct EVE pattern was observed. Across three scaffolds, 37 loci showed homology to 31 CjLGDV genes, with 35 concentrated on one scaffold in the length of about 100 kb (NW_020229367.1). These loci were confirmed as endogenous fragments based on several lines of evidence: the presence of identical sequences in another *Camponotus floridanus* genome assembly, comparable sequencing depth to host BUSCO scaffolds, and the occurrence of multiple premature stop codons and transposable elements within these regions. RNA-seq analyses across multiple datasets did not detect any transcriptional activity in the EVE regions in *Camponotus floridanus*.

Gene synteny analysis of five representative EVEs containing multiple viral gene homologs revealed that all the EVEs had some degree of gene collinearity with CjLGDV (Supplementary Fig. S11A), supporting the evolutionary linkage between CjLGDV and the ant hosts. The 100 kb long region within the scaffold (NW_020229367.1) in *Camponotus floridanus* genome showed high synteny with the CjLGDV genome (Supplementary Fig. S11B), suggesting that this scaffold was very likely derived from a virus closely related to CjLGDV.

### Ant EVEs and ants have engaged in long-term coevolution

To investigate the selective pressures acting on ant EVEs, we analyzed the ratio of nonsynonymous to synonymous substitution rates (*d_N_*/*d_S_*). The results revealed that most CjLGDV homologs in ant EVEs have undergone strong purifying selection (*d_N_*/*d_S_* < 1; Supplementary Fig. S12, Supplementary Table S5). By searching available RNA-seq data, it was shown that at least 135 EVEs in 13 ant species were transcriptionally active (Supplementary Table S5), of which 67 exhibited *d_N_*/*d_S_* < 1, suggesting evolutionary constraints on these EVEs and their functional importance.

As the *odv-e66* homologs in EVEs were conserved in most ant species, including 26 ant species across all five subfamilies (Fig. 5), we performed phylogenetic analysis using *odv-e66* homologs from bacteria, viruses and ant EVEs to examine their evolutionary relationships. The *odv-e66* homologs in viruses clustered in accordance to their taxonomic classifications (Fig. 6A). Most *odv-e66* homologs in ant EVEs grouped together, forming a phylogenetic topology that mirrored the evolutionary relationships of their ant hosts (Fig. 6B). Similar co-phylogenetic pattern was observed in other viral homologs in EVEs (Supplementary Fig. S9), suggesting that the integration of EVEs have followed the evolutionary history of their hosts. However, some ant species, such as *Camponotus floridanus*, *Formica aserva*, *Lasius niger*, contained *odv-e66* homologs that phylogenetically clustered with CjLGDV and CjLGDV-like, which indicated ongoing interactions between viruses and their host genomes or multiple integration events might have occurred over evolutionary time.

**Fig. 6.**
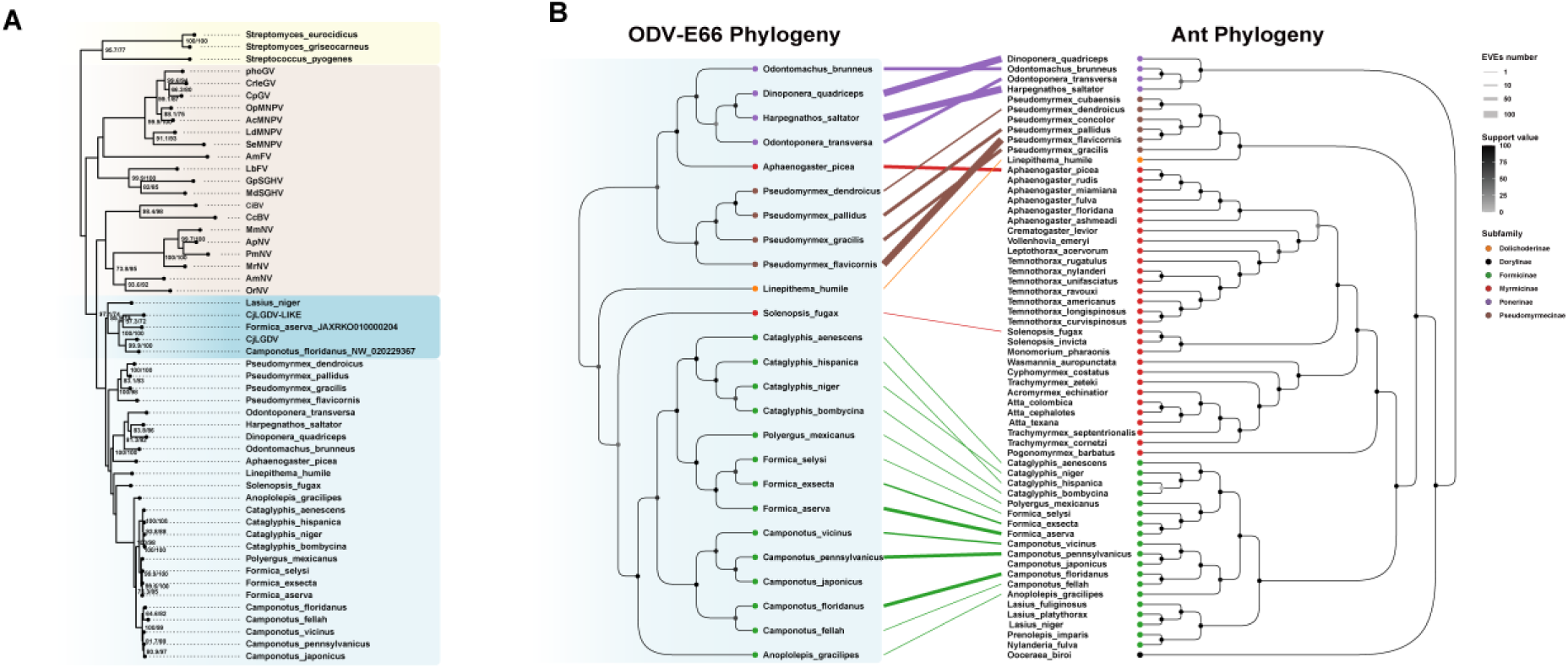
Relationship between ant EVEs and their ant hosts. (A) The phylogenetic tree was constructed using maximum likelihood (ML) inference based on the codon alignments of *odv-e66* homologs in bacterial, viruses and EVEs. Node support values are indicated as SH-aLRT support (%) / Ultrafast bootstrap (%). (B) Tanglegram shows the relationship between ant *odv-e66* homologs and their ant hosts. Labels and connecting lines are colored by ant subfamilies. The thickness of connecting lines represented the number of EVEs. Ant phylogeny was inferred from the concatenated amino acid sequences of 106 BUSCO genes.

Using the relaxed clock model, we calculated the divergence times based on the codon alignments of *odv-e66* homologs, with calibration points including the estimated time to the most recent common ancestor (TMRCA) of Chelonus inanitus bracovirus (CiBV) and Cotesia congregata bracovirus (CcBV) (Thézé et al. 2011), as well as the divergence time of Formicinae ants (Blanchard and Moreau 2017). It is estimated the TMRCA of all lefaviruses *odv-e66* homologs is approximately 456 million years ago (Mya) (Fig. 7). The TMRCAs of lefaviruses and bacteria *odv-e66* homologs, including those from *Streptococcus pyogenes*, *Streptomyces eurocidicus* and *Streptomyces griseocarneus*, is estimated at ∼535 Mya. The TMRCA of ant EVEs and ant viruses is estimated to be ∼ 213 Mya. For CjLGDV and CjLGDV-like, the TMRCA is estimated to be ∼125 Mya. Notably, the TMRCA of ant EVEs that mirror the evolutionary relationships of their ant hosts is estimated at ∼187 Mya. The divergence times of these ant EVEs is close to the divergence times of their respective host species, further supporting a co-evolutionary relationship.

**Fig. 7.**
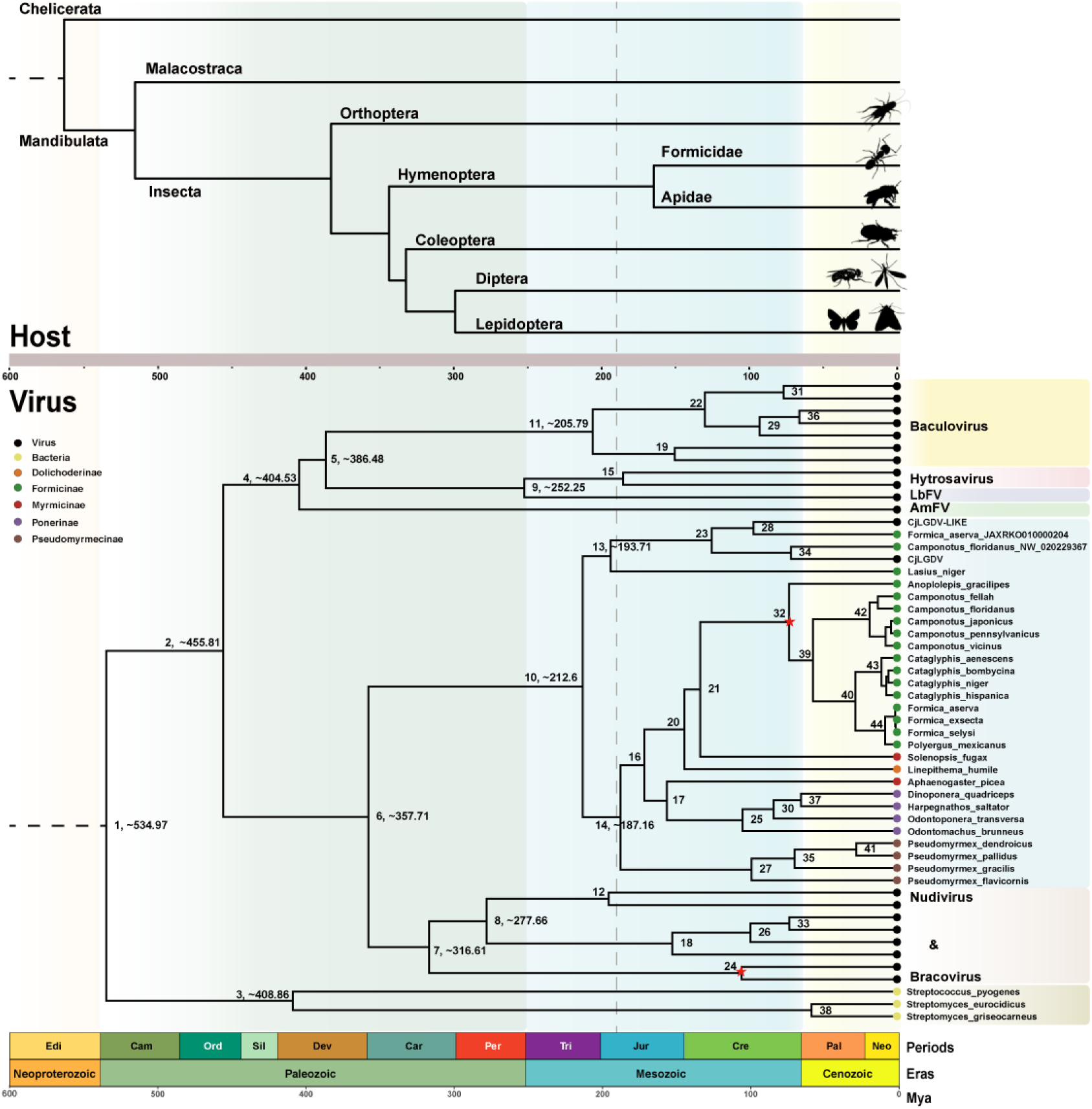
Bayesian timetree of the *odv-e66* homologs in bacteria, viruses and EVEs. Calibrated nodes are marked with red stars. Age estimates and their 95% HPD intervals for the corresponding nodes are provided in Supplementary Table S6. The host time tree was obtained from Timetree.

## Discussion

Substantial progress has been made in elucidating the viral landscape in invertebrates (Shi et al. 2016; Williams et al. 2017), with various insects identified as hosts to large DNA viruses (Gauthier et al. 2015; Lepetit et al. 2016; Abd-Alla et al. 2008; Bézier et al. 2014; Wang et al. 2007). However, few DNA viruses infecting ants have been characterized so far (Valles et al. 2013; Baty et al. 2020; Flynn and Moreau 2024).

This study presents the first fully sequenced ant dsDNA virus from ants, which is tentatively named as Camponotus japonicus labial gland disease virus (CjLGDV). The observation of abundant viral particles and high ratio of DNA sequencing reads (41.20% of CjLGDV vs. 48.90% of ant host) in the enlarged labial gland suggest that CjLGDV may be associated with ‘labial gland disease’. The observation of elongated nucleocapsids stacked in the nucleus, along with coiled and uncoiled nucleocapsids with envelopes in the cytoplasm, suggests a lifecycle of CjLGDV similar to that of most DNA viruses that replicate in the nucleus and mature in the cytoplasm. By sequence assembly, it is revealed that CjLGDV contains a 142 kb circular dsDNA genome. By data mining in public databases, we also identified a 104kb CjLGDV-like scaffold in *Anoplolepis gracilipes*, which showed high gene synteny with the CjLGDV genome. The identification of CjLGDV and CjLGDV-like scaffold suggests a possibility of the existence of a new clade of DNA viruses infecting ants.

CjLGDV and CjLGDV-like exhibit similarities to arthropod large DNA viruses in terms of GC content, genome size, coding density and the existence of repeat sequences. The identification of NALDVs gene homologs in CjLGDV and CjLGDV-like genomes supports their association with NALDVs. Additionally, the presence of *pif* genes homologs and three DNA-directed RNA polymerase subunits (*lef4*, *lef8*, and *lef9*) suggests a linkage of the two viruses to *Lefavirales*. Phylogenetic analysis clusters CjLGDV and CjLGDV-like together as a novel entity within the *Lefavirales*, forming a monophyletic clade with unclassified bee viruses AmFV and AmFLV, while LbFV and hytrosaviruses form a closely related sister clade. This unique placement highlights the distinct evolutionary lineage of CjLGDV and CjLGDV-like, differentiating them from baculoviruses and nudiviruses, and also suggesting potential functional and evolutionary links of the two novel ant viruses to AmFV, AmFLV, LbFV and hytrosaviruses. The close relationship of CjLGDV and CjLGDV-like to AmFV and AmFLV raises intriguing questions about their evolutionary trajectory. Considering the intersecting ecological niches and interspecies interactions between ants and bees, researches have identified ants as plausible vectors and reservoirs for pathogens affecting bee populations (Payne et al. 2020; Schläppi et al. 2020; Celle et al. 2008; Levitt et al. 2013; Lin et al. 2020; Dobelmann et al. 2023). It is reasonable to hypothesize that these viruses may have evolved close associations due to their host interactions. The enlarged labial gland observed in *Camponotus japonicus* is reminiscent of the salivary gland hypertrophy seen in flies infected with hytrosaviruses (Zhang et al. 2023; Kariithi et al. 2017), potentially indicating a shared pathological mechanism with a common evolutionary origin. Hytrosavirus-like viral genome fragments have been detected in *Gigantiops destructor* (Flynn and Moreau 2024), indicating the possibility of *Hytrosaviridae* viral infection within ants and a possible connection between CjLGDV and hytrosaviruses.

The LbFV is associated with superparasitism behavior of *Leptopilina boulardi* (Varaldi et al. 2006; Patot et al. 2009), and the virus infection may contribute adaptive genes to parasitic wasps (Burke et al. 2021; Di Giovanni et al. 2020). Viral manipulation of host behavior represents a sophisticated evolutionary strategy that optimizes transmission conditions and enhances survival, illustrating the complex interplay between virus and host (Varaldi et al. 2003; Dheilly et al. 2015). Further research is required to determine whether CjLGDV induces behavioral modifications in ants, which could deepen our understanding of host-pathogen dynamics.

EVEs are the result of chromosomal integration of viral genes in the host germline cells. Non-retroviral EVEs represent rare remnants of ancient viral infections. Study on these EVEs can offer valuable insights into viral host range, ancestral viral diversity, and the timing of viral evolutionary events. In this study, an unprecedented number of EVEs which were more closely related to CjLGDV and CjLGDV-like rather than AmFV or any other known viruses, were detected in the genomes of various ant species spanning five subfamilies (*Myrmicinae*, *Formicinae*, *Ponerinae*, *Dolichoderinae* and *Pseudomyrmicinae*). The evolutionary history of these EVEs largely follows host genetic inheritance patterns. The TMRCA of all ant EVEs and CjLGDV was estimated at ∼213 Mya, whereas the TMRCA of ant EVEs that reflected the evolutionary relationships of their ant hosts was ∼187 Mya—closely aligning with the divergence time of the family *Formicidae*. Based on the broad detection of CjLGDV-related EVEs and the co-phylogenetic patterns observed between these EVEs and their ant hosts, we speculate that the ancestor of these ant species have served as hosts to the ancestor of CjLGDV. Considering the low probability of non-retroviral endogenization events occurring in the germline, the presence of EVEs strongly indicates prolonged and frequent interactions between the ancestral viruses and their hosts. The discovery of CjLGDV in *Camponotus japonicus* and CjLGDV-like in *Anoplolepis gracilipes*, along with reports of “labial gland disease” in various ant species, implies that such interactions persist in their descendants. Some EVEs identified in this study exhibit high gene collinearity and close phylogenetic relationships with CjLGDV, indicating the occurrance of some recent integration events or the integration from more closely related viruses. All these findings strongly support the hypothesis of long-term and frequent interactions between CjLGDV and ants. Additional efforts on the direct detection and characterization of viruses from ants are necessary to conclusively establish the natural host range of CjLGDV and to elucidate the relationship between CjLGDV and CjLGDV-related EVEs.

Studies have shown that EVEs can serve as reservoirs of immune memory in hosts and may function in antiviral defense (Whitfield et al. 2017; Ligoxygakis 2017; Kloc et al. 2024; Lanz-Mendoza and Contreras-Garduño 2022). In some parasitic wasps, endogenous viral elements have been discovered to produce “virus-like structures” (VLS) in females’ reproductive organs, which can deliver immunosuppressive DNA or proteins to modulate the immune system of their hosts (Di Giovanni et al. 2020; Bézier et al. 2009; Pichon et al. 2015). In our study, viral homologs in CjLGDV-related EVEs are found to be under strong purifying selection, and the transcription of some EVEs in some ant species can be spotted from the public available databases. These findings suggest the potential domestication of CjLGDV-related EVEs for functional roles. One such gene, *odv-e66* which encodes a chondroitinase associated with viral infectivity, has been hypothesized to be a key factor in virus-host adaptation (Gauthier et al. 2021). Here we reveal that homologs of *odv-e66* are widely distributed in diverse ant species, and some of them are transcriptionally active. Whether ODV-E66 plays a functional role and contributes to ant evolution, it remains to be a question that requires further investigation.

Taken together, we identified a novel dsDNA virus CjLGDV, which is potentially responsible for labial gland hypertrophy in *Camponotus japonicus*. Phylogenetic analyses demonstrate that CjLGDV belongs to a new clade in the order *Lefavirales*, class *Naldaviricetes*. The detection of CjLGDV-related endogenous viral elements in various ant genomes provides evidence that ants have been the host to CjLGDV or its ancestors with a long history. These findings expand the known lineage of NALDVs, broaden our understanding of their host range, and reveal a long and frequent interactions between CjLGDV and ant host.

## Materials and Methods

### Ant collection

Colonies of *Camponotus japonicus*, i<u>n</u> which minor workers were observed to have enlarged labial glands (Zhang et al. 2023), were collected from Yangling, Shaanxi Province. The ants were refrigerated at −20°C for 10 minutes to reduce their activity, after which their labial glands were dissected in Ringer’s physiological solution under an Olympus SZ51 microscope.

### Electron microscopy

Labial glands were initially fixed in 2.5% cold glutaraldehyde for at least 12 h, then washed five times with PBS buffer (0.1 M, pH 7.2). Post-fixation treatment was performed in 1% osmium tetroxide for 2 hours, followed by five washes with PBS. Samples were dehydrated through a graded ethanol series (30%, 50%, 70%, 80%, 90%, and 100%) and 100% acetone. The tissue was infiltrated with LR-White resin and acetone mixtures (1:3 for 2 hours, 1:1 for 5 hours, and 3:1 for 12 hours), then embedded in pure LR-White resin and polymerized over a period of 72 hours. The thin sections were obtained using a Leica EM UC7 ultramicrotome (Hitachi, Tokyo, Japan), and double-stained using uranyl acetate for 20 minutes followed by lead citrate for 10 minutes to enhance contrast. Observation of the stained sections wa<u>s</u> conducted using a Tecnai G2 Spirit Bio Twin electron microscope (FEI, Chech, USA).

### Genomic DNA preparation and sequencing

Normal labial glands (LG) and enlarged labial glands (ENLG) were obtained from ants in the same colony. The genomic DNA was extracted from the entire labial glands with a genomic DNA extraction kit (BioTeke, Beijing, China) following the manufacturer’s protocol. The quantity and quality of extracted DNAs were measured using a NanoDrop ND-1000 spectrophotometer (Thermo Fisher Scientific, Waltham, MA, USA) and agarose gel electrophoresis, respectively. The extracted DNA was processed to construct metagenome shotgun sequencing libraries with insert size of 400 bp by using Illumina TruSeq Nano DNA LT Library Preparation Kit. The library was sequenced on Illumina HiSeq X-ten platform (Illumina, USA) with PE150 strategy by Personal Biotechnology Co., Ltd. (Shanghai, China).

### Viral genome assembly

Paired-end sequencing reads were quality-filtered and trimmed using Cutadapt v4.4 (Martin 2011). Genomes of three species closely related to *Camponotus japonicus*—*Camponotus floridanus*, *Camponotus pennsylvanicus* and *Camponotus vicinus*—were merged into a composite reference genome. Sequencing data from the LG and ENLG groups were aligned to the reference using Bowtie2 v2.5 (Langmead and Salzberg 2012). Reads that aligned to reference genome were regarded as the host sequences, and the remaining reads were utilized for viral genome assembly using Haploflow (Fritz et al. 2021). The quality of the assembly was evaluated using QUAST v5.2 (Gurevich et al. 2013). All reads used for genome assembly were mapped to the assembled contigs using Bowtie2, with coverage determined by bedtools v2.31 (Quinlan and Hall 2010). Low abundance contigs were filtered out, and overlapping ones were merged. To resolve gaps and ambiguous regions between contigs, PCR was performed using primers listed in Supplementary Table S1, and the sequences of the amplified fragments were determined by Sanger sequencing.

### Sequence analyses and genome annotation

Whole-genome similarities were evaluated by employing TBLASTX (Altschul et al. 1997, 2005) against a set of representative invertebrate DNA virus genomes. Putative open reading frames (ORFs) were identified using ORF finder and Prodigal v2.6 (Hyatt et al. 2010). ORFs were named based on their homologs or genomic location. BLASTP was used to identify ORF similarities against the NCBI nonredundant protein database. Domain identification within ORFs was performed using the NCBI Conserved Domain Search (Wang et al. 2023) and HMMER-search against both public databases (CDD, PFAM) and local databases. The local databases were built using homologous sequences from nuclear arthropod large DNA viruses, which were aligned using MAFFT v7.5 (Nakamura et al. 2018) and converted into HMMs using hmmbuild. Protein structure alignment and homologous structure searches were conducted using Foldseek (van Kempen et al. 2024). The sequence coding density (CD) was calculated as the ratio of the base number of all ORFs to the base number of the genome. Tandem direct repeats and imperfect palindromic motifs were identified using the etandem (Rice et al. 2000) and MEME suite (Bailey et al. 2006), respectively, with 100 score cutoff. The virus genome was graphically represented in a circular diagram using CGView (Stothard and Wishart 2005).

### Phylogenetic analysis

The phylogenetic position of CjLGDV within *Naldaviricetes* was inferred by maximum likelihood tree based on 12 conserved genes. Sequence accession numbers for the conserved genes used in the analysis are provided in Supplementary Table S4. Specifically, amino acid sequences were aligned using MAFFT with the E-INS-I mode. These alignments were subsequently trimmed by trimAl v1.4 (Capella-Gutiérrez et al. 2009) and concatenated into a single protein alignment by SequenceMatrix (Vaidya et al. 2011). Phylogenetic trees were then constructed using the maximum-likelihood method implemented in IQ-TREE v2.2 (Minh et al. 2020). ModelFinder (Kalyaanamoorthy et al. 2017) was employed within IQ-TREE to identify the best models for each partition. White spot syndrome virus (WSSV) was selected as the outgroup. Node supports in the ML trees were determined using Ultra-fast bootstrap (Hoang et al. 2018) and SH-aLRT (options -bb 1000 and -alrt 1000). The patristic distances within and between viral families were performed by the ape R package (Paradis and Schliep 2019).

### Identification of endogenous viral elements

A BLAST-based approach complemented by systematic phylogenetic clade validations was utilized to identify endogenous viral sequences. Putative protein sequences of CjLGDV were subjected to a TBLASTN search against a database comprising 59 ant genomes. Information of these genomes are provided in Supplementary Table S7. Only hits with e-value smaller than 1e-5 and more than 20% sequence alignment coverage were maintained. Adjacent hits within distance of 10 bp were merged into a single entity. Hits were then subjected to a BLASTX search against the NCBI nonredundant protein database. Sequences that clearly align with viral sequences were identified as potential EVEs. Several criteria were utilized to evaluate the endogenous characteristics of candidate EVEs, including the presence of transposable elements, insect genes, premature stop codons and sequencing depth. Transposable elements on the scaffolds were predicted using EDTA v2.0 (Ou et al. 2019), while genes were predicted using AUGUSTUS v3.5 (Stanke et al. 2008). Taxonomic assignments were performed on sequence similarity with the Uniprot/Swissprot database by BLASTP. Only genes assigned to insects were retained. Details on these EVE sequences are available in Supplementary Table S5. To perform phylogenetic analyses, EVEs were aligned with related homologs from CjLGDV and NALDVs using MAFFT v7.5 and subsequently refined with trimAl v1.4. Phylogenetic trees were constructed using the maximum-likelihood method implemented in IQ-TREE v2.2. To assess potential functional constraints on the EVEs, the ratio of nonsynonymous substitution rate (*d_N_*) to synonymous substitution rate (*d_S_*) was estimated using codeml on PAML package (Yang 2007).

### EVE-Host Evolutionary Association Analyses

To investigate the co-phylogenetic relationships of EVEs and their hosts, we compared the topological structure between the host and EVEs phylogenies. Phylogenetic analysis of ODV-E66 homologs from virus and ants was conducted. The ant phylogeny was inferred from the concatenated amino acid sequences of 106 BUSCO genes using the maximum-likelihood method implemented in IQ-TREE v2.2. The relationship between ODV-E66 and ant phylogeny was visualized in a tanglegram by the ape v5.0 R package (Paradis and Schliep 2019).

### Molecular Dating Analysis

Codon alignments for *odv-e66* homologs, obtained using MACSE2 (Ranwez et al. 2018), were used for molecular dating analysis. The phylogenetic tree was constructed using the Bayesian inference method implemented in BEAST2 (Bouckaert et al. 2019). The analysis employed the optimized relaxed clock (ORC) (Douglas et al. 2021), and Monte Carlo Markov Chains (MCMC) for 200 million generations. The divergence times of CiBV and CcBV (Thézé et al. 2011; Whitfield 2002), as well as the divergence time of the Formicinae ants (Blanchard and Moreau 2017) were used as calibration points. The MCMC runs were diagnosed using Tracer (Rambaut et al. 2018) ensuring that all effective sampling sizes were larger than 200.

### Data access

The data underlying this article are available in the article and in its online supplementary material. The CjLGDV genome sequence has been deposited in the GenBank Nucleotide Database under accession number PP861292. The DNA sequencing data generated in this study has been deposited in the NCBI SRA database under accession number PRJNA1119426 and is publicly available as of the date of publication.

## Acknowledgments

We are grateful to Kerang Huang and Zhen Wang (Life Science Core Services Facility, Northwest A&F University, China) for assistance in transmission electron microscopy.

This work was supported by the National Natural Science Foundation of China (Grants No. 32071490) and the National Natural Science Foundation of China (Grants No. 32371566).

## References

Abd-Alla AMM, Cousserans F, Parker AG, Jehle JA, Parker NJ, Vlak JM, Robinson AS, Bergoin M. 2008. Genome analysis of a Glossina pallidipes salivary gland hypertrophy virus reveals a novel, large, double-stranded circular DNA virus. J Virol 82: 4595–4611.

Altschul SF, Madden TL, Schäffer AA, Zhang J, Zhang Z, Miller W, Lipman DJ. 1997. Gapped BLAST and PSI-BLAST: a new generation of protein database search programs. Nucleic Acids Research 25: 3389–3402.

Altschul SF, Wootton JC, Gertz EM, Agarwala R, Morgulis A, Schäffer AA, Yu Y-K. 2005. Protein database searches using compositionally adjusted substitution matrices. FEBS J 272: 5101–5109.

BabéLM, Craik CS. 1997. Viral Proteases: Evolution of Diverse Structural Motifs to Optimize Function. Cell 91: 427–430.

Bailey TL, Williams N, Misleh C, Li WW. 2006. MEME: discovering and analyzing DNA and protein sequence motifs. Nucleic Acids Res 34: W369–373.

Baty J, Bulgarella M, Dobelmann J, Felden A, Lester PJ. 2020. Viruses and their effects in ants (Hymenoptera: Formicidae). Myrmecological News 30: 213–228.

Bazan JF, Fletterick RJ. 1988. Viral cysteine proteases are homologous to the trypsin-like family of serine proteases: structural and functional implications. Proc Natl Acad Sci U S A 85: 7872–7876.

Bézier A, Herbinière J, Lanzrein B, Drezen J-M. 2009. Polydnavirus hidden face: The genes producing virus particles of parasitic wasps. Journal of Invertebrate Pathology 101: 194– 203.

Bézier A, ThézéJ, Gavory F, Gaillard J, Poulain J, Drezen J-M, Herniou EA. 2014. The Genome of the Nucleopolyhedrosis-Causing Virus from Tipula oleracea Sheds New Light on the Nudiviridae Family. J Virol 89: 3008–3025.

Blanchard BD, Moreau CS. 2017. Defensive traits exhibit an evolutionary trade-off and drive diversification in ants. Evolution 71: 315–328.

Bouckaert R, Vaughan TG, Barido-Sottani J, Duchêne S, Fourment M, Gavryushkina A, Heled J, Jones G, Kühnert D, De Maio N, et al. 2019. BEAST 2.5: An advanced software platform for Bayesian evolutionary analysis. PLoS Comput Biol 15: e1006650.

Briese T, Kapoor A, Mishra N, Jain K, Kumar A, Jabado OJ, Lipkin WI. 2015. Virome Capture Sequencing Enables Sensitive Viral Diagnosis and Comprehensive Virome Analysis. mBio 6: e01491–15.

Burke GR, Hines HM, Sharanowski BJ. 2021. The Presence of Ancient Core Genes Reveals Endogenization from Diverse Viral Ancestors in Parasitoid Wasps. Genome Biology and Evolution 13: evab105.

Burke GR, Thomas SA, Eum JH, Strand MR. 2013. Mutualistic Polydnaviruses Share Essential Replication Gene Functions with Pathogenic Ancestors. PLoS Pathog 9: e1003348.

Capella-Gutiérrez S, Silla-Martínez JM, Gabaldón T. 2009. trimAl: a tool for automated alignment trimming in large-scale phylogenetic analyses. Bioinformatics 25: 1972–1973.

Celle O, Blanchard P, Olivier V, Schurr F, Cougoule N, Faucon J-P, Ribière M. 2008. Detection of Chronic bee paralysis virus (CBPV) genome and its replicative RNA form in various hosts and possible ways of spread. Virus Res 133: 280–284.

Dheilly NM, Maure F, Ravallec M, Galinier R, Doyon J, Duval D, Leger L, Volkoff A-N, Missé D, Nidelet S, et al. 2015. Who is the puppet master? Replication of a parasitic wasp-associated virus correlates with host behaviour manipulation. Proc Biol Sci 282: 20142773.

Di Giovanni D, Lepetit D, Guinet B, Bennetot B, Boulesteix M, CoutéY, Bouchez O, Ravallec M, Varaldi J. 2020. A Behavior-Manipulating Virus Relative as a Source of Adaptive Genes for Drosophila Parasitoids. Mol Biol Evol 37: 2791–2807.

Dobelmann J, Felden A, Lester PJ. 2023. An invasive ant increases deformed wing virus loads in honey bees. Biol Lett 19: 20220416.

Douglas J, Zhang R, Bouckaert R. 2021. Adaptive dating and fast proposals: Revisiting the phylogenetic relaxed clock model. PLOS Computational Biology 17: e1008322.

Dubrez-Daloz L, Dupoux A, Cartier J. 2008. IAPs: more than just inhibitors of apoptosis proteins. Cell Cycle 7: 1036–1046.

Elton ETG. 1991. Labial gland disease in the genusFormica (Formicidae, Hymenoptera). Ins Soc 38: 91–93.

Flynn PJ, Moreau CS. 2019. Assessing the Diversity of Endogenous Viruses Throughout Ant Genomes. Front Microbiol 10: 1139.

Flynn PJ, Moreau CS. 2024. Viral diversity and co-evolutionary dynamics across the ant phylogeny. Mol Ecol e17519.

Fritz A, Bremges A, Deng Z-L, Lesker TR, Götting J, Ganzenmueller T, Sczyrba A, Dilthey A, Klawonn F, McHardy AC. 2021. Haploflow: strain-resolved de novo assembly of viral genomes. Genome Biol 22: 212.

Garcia-Maruniak A, Maruniak JE, Farmerie W, Boucias DG. 2008. Sequence analysis of a non-classified, non-occluded DNA virus that causes salivary gland hypertrophy of Musca domestica, MdSGHV. Virology 377: 184–196.

Gauthier J, Boulain H, van Vugt JJFA, Baudry L, Persyn E, Aury J-M, Noel B, Bretaudeau A, Legeai F, Warris S, et al. 2021. Chromosomal scale assembly of parasitic wasp genome reveals symbiotic virus colonization. Commun Biol 4: 104.

Gauthier L, Cornman S, Hartmann U, Cousserans F, Evans JD, de Miranda JR, Neumann P. 2015. The Apis mellifera Filamentous Virus Genome. Viruses 7: 3798–3815.

Gilbert C, Belliardo C. 2022. The diversity of endogenous viral elements in insects. Current Opinion in Insect Science 49: 48–55.

Guarino LA, Xu B, Jin J, Dong W. 1998. A virus-encoded RNA polymerase purified from baculovirus-infected cells. J Virol 72: 7985–7991.

Guinet B, Lepetit D, Charlat S, Buhl PN, Notton DG, Cruaud A, Rasplus J-Y, Stigenberg J, de Vienne DM, Boussau B, et al. 2023. Endoparasitoid lifestyle promotes endogenization and domestication of dsDNA viruses eds. A. Rokas and C.R. Landry. eLife 12: e85993.

Gurevich A, Saveliev V, Vyahhi N, Tesler G. 2013. QUAST: quality assessment tool for genome assemblies. Bioinformatics 29: 1072–1075.

Hoang DT, Chernomor O, von Haeseler A, Minh BQ, Vinh LS. 2018. UFBoot2: Improving the Ultrafast Bootstrap Approximation. Mol Biol Evol 35: 518–522.

Hou D, Kuang W, Luo S, Zhang F, Zhou F, Chen T, Zhang Y, Wang H, Hu Z, Deng F, et al. 2019. Baculovirus ODV-E66 degrades larval peritrophic membrane to facilitate baculovirus oral infection. Virology 537: 157–164.

Hyatt D, Chen G-L, Locascio PF, Land ML, Larimer FW, Hauser LJ. 2010. Prodigal: prokaryotic gene recognition and translation initiation site identification. BMC Bioinformatics 11: 119.

Jin J, Dong W, Guarino LA. 1998. The LEF-4 subunit of baculovirus RNA polymerase has RNA 5’-triphosphatase and ATPase activities. J Virol 72: 10011–10019.

Kadlečková D, Saláková M, Erban T, Tachezy R. 2024. Discovery and characterization of novel DNA viruses in Apis mellifera: expanding the honey bee virome through metagenomic analysis. Msystems 0: e00088–24.

Kalyaanamoorthy S, Minh BQ, Wong TKF, von Haeseler A, Jermiin LS. 2017. ModelFinder: fast model selection for accurate phylogenetic estimates. Nat Methods 14: 587–589.

Kariithi HM, Meki IK, Boucias DG, Abd-Alla AM. 2017. Hytrosaviruses: current status and perspective. Curr Opin Insect Sci 22: 71–78.

Kawato S, Shitara A, Wang Y, Nozaki R, Kondo H, Hirono I. 2019. Crustacean Genome Exploration Reveals the Evolutionary Origin of White Spot Syndrome Virus. J Virol 93: e01144–18.

Ke J, Wang J, Deng R, Lin L, Jinlong B, Yaoping L, Wang X. 2011. Characterization of AcMNPV with a deletion of ac69 gene. Microbiology Research 2: e4.

Kloc M, Halasa M, Kubiak JZ, Ghobrial RM. 2024. Invertebrate Immunity, Natural Transplantation Immunity, Somatic and Germ Cell Parasitism, and Transposon Defense. Int J Mol Sci 25: 1072.

Kool M, Ahrens CH, Vlak JM, Rohrmann GF. 1995. Replication of baculovirus DNA. J Gen Virol 76 ( **Pt 9**): 2103–2118.

Laciny A. 2021. Among the shapeshifters: parasite-induced morphologies in ants (Hymenoptera, Formicidae) and their relevance within the EcoEvoDevo framework. Evodevo 12: 2.

Langmead B, Salzberg SL. 2012. Fast gapped-read alignment with Bowtie 2. Nat Methods 9: 357– 359.

Lanz-Mendoza H, Contreras-Garduño J. 2022. Innate immune memory in invertebrates: Concept and potential mechanisms. Dev Comp Immunol 127: 104285.

Leisy DJ, Rohrmann GF. 1993. Characterization of the replication of plasmids containing hr sequences in baculovirus-infected Spodoptera frugiperda cells. Virology 196: 722–730.

Lepetit D, Gillet B, Hughes S, Kraaijeveld K, Varaldi J. 2016. Genome Sequencing of the Behavior Manipulating Virus LbFV Reveals a Possible New Virus Family. Genome Biology and Evolution 8: 3718–3739.

Levitt AL, Singh R, Cox-Foster DL, Rajotte E, Hoover K, Ostiguy N, Holmes EC. 2013. Cross-species transmission of honey bee viruses in associated arthropods. Virus Res 176: 232– 240.

Li Q, Lian Y, Zhang K, Chen J, Chen L, Wu J, Zhang Y, Chen M, Zhang W, Lu M, et al. 2024. Virome of red imported fire ants by metagenomic analysis in Guangdong, southern China. Front Microbiol 15: 1479934.

Li Y, Hu L, Chen T, Chang M, Deng F, Hu Z, Wang H, Wang M. 2020. Host AAA+ ATPase TER94 Plays Critical Roles in Building the Baculovirus Viral Replication Factory and Virion Morphogenesis. J Virol 94: e01674–19.

Ligoxygakis P. 2017. Immunity: Insect Immune Memory Goes Viral. Current Biology 27: R1218–R1220.

Lin B, Averett WF, Pritchard DG. 1997. Identification of a Histidine Residue Essential for Enzymatic Activity of Group B Streptococcal Hyaluronate Lyase. Biochemical and Biophysical Research Communications 231: 379–382.

Lin C-Y, Lee C-C, Nai Y-S, Hsu H-W, Lee C-Y, Tsuji K, Yang C-CS. 2020. Deformed Wing Virus in Two Widespread Invasive Ants: Geographical Distribution, Prevalence, and Phylogeny. Viruses 12: 1309.

Martin M. 2011. Cutadapt removes adapter sequences from high-throughput sequencing reads. EMBnet.journal 17: 10–12.

Minh BQ, Schmidt HA, Chernomor O, Schrempf D, Woodhams MD, von Haeseler A, Lanfear R. 2020. IQ-TREE 2: New Models and Efficient Methods for Phylogenetic Inference in the Genomic Era. Mol Biol Evol 37: 1530–1534.

Nakamura T, Yamada KD, Tomii K, Katoh K. 2018. Parallelization of MAFFT for large-scale multiple sequence alignments. Bioinformatics 34: 2490–2492.

Ogura T, Wilkinson AJ. 2001. AAA+ superfamily ATPases: common structure–diverse function. Genes to Cells 6: 575–597.

Ou S, Su W, Liao Y, Chougule K, Agda JRA, Hellinga AJ, Lugo CSB, Elliott TA, Ware D, Peterson T, et al. 2019. Benchmarking transposable element annotation methods for creation of a streamlined, comprehensive pipeline. Genome Biol 20: 275.

Paradis E, Schliep K. 2019. ape 5.0: an environment for modern phylogenetics and evolutionary analyses in R. Bioinformatics 35: 526–528.

Patot S, Lepetit D, Charif D, Varaldi J, Fleury F. 2009. Molecular Detection, Penetrance, and Transmission of an Inherited Virus Responsible for Behavioral Manipulation of an Insect Parasitoid. Applied and Environmental Microbiology 75: 703–710.

Payne AN, Shepherd TF, Rangel J. 2020. The detection of honey bee (Apis mellifera)-associated viruses in ants. Sci Rep-uk 10: 2923.

Pichon A, Bézier A, Urbach S, Aury J-M, Jouan V, Ravallec M, Guy J, Cousserans F, ThézéJ, Gauthier J, et al. 2015. Recurrent DNA virus domestication leading to different parasite virulence strategies. Science Advances 1: e1501150.

Quinlan AR, Hall IM. 2010. BEDTools: a flexible suite of utilities for comparing genomic features. Bioinformatics 26: 841–842.

Rambaut A, Drummond AJ, Xie D, Baele G, Suchard MA. 2018. Posterior Summarization in Bayesian Phylogenetics Using Tracer 1.7. Syst Biol 67: 901–904.

Ranwez V, Douzery EJP, Cambon C, Chantret N, Delsuc F. 2018. MACSE v2: Toolkit for the Alignment of Coding Sequences Accounting for Frameshifts and Stop Codons. Molecular Biology and Evolution 35: 2582–2584.

Rascovan N, Duraisamy R, Desnues C. 2016. Metagenomics and the Human Virome in Asymptomatic Individuals. Annual Review of Microbiology 70: 125–141.

Rice P, Longden I, Bleasby A. 2000. EMBOSS: the European Molecular Biology Open Software Suite. Trends Genet 16: 276–277.

Rodrigues DT, Peterson L, de Oliveira LB, Sosa-Gómez DR, Ribeiro BM, Ardisson-Araújo DMP. 2020. Characterization of a novel alphabaculovirus isolated from the Southern armyworm, *Spodoptera eridania* (Cramer, 1782) (Lepidoptera: Noctuidae) and the evolution of *odv-e66*, a bacterium-acquired baculoviral chondroitinase gene. Genomics 112: 3903–3914.

Schläppi D, Chejanovsky N, Yañez O, Neumann P. 2020. Foodborne Transmission and Clinical Symptoms of Honey Bee Viruses in Ants Lasius spp. Viruses 12: 321.

Shi M, Lin X-D, Tian J-H, Chen L-J, Chen X, Li C-X, Qin X-C, Li J, Cao J-P, Eden J-S, et al. 2016. Redefining the invertebrate RNA virosphere. Nature 540: 539–543.

Snider J, Thibault G, Houry WA. 2008. The AAA+ superfamily of functionally diverse proteins. Genome Biology 9: 216.

Stanke M, Diekhans M, Baertsch R, Haussler D. 2008. Using native and syntenically mapped cDNA alignments to improve de novo gene finding. Bioinformatics 24: 637–644.

Stothard P, Wishart DS. 2005. Circular genome visualization and exploration using CGView. Bioinformatics 21: 537–539.

Takemoto Y, Mitsuhashi W, Murakami R, Konishi H, Miyamoto K. 2008. The N-Terminal Region of an Entomopoxvirus Fusolin Is Essential for the Enhancement of Peroral Infection, whereas the C-Terminal Region Is Eliminated in Digestive Juice. J Virol 82: 12406–12415.

Theilmann DA, Stewart S. 1992. Tandemly repeated sequence at the 3’ end of the IE-2 gene of the baculovirus Orgyia pseudotsugata multicapsid nuclear polyhedrosis virus is an enhancer element. Virology 187: 97–106.

ThézéJ, Bézier A, Periquet G, Drezen J-M, Herniou EA. 2011. Paleozoic origin of insect large dsDNA viruses. Proceedings of the National Academy of Sciences 108: 15931–15935.

Vaidya G, Lohman DJ, Meier R. 2011. SequenceMatrix: concatenation software for the fast assembly of multi-gene datasets with character set and codon information. Cladistics 27: 171–180.

Valles SM. 2024. Effect of Solenopsis invicta virus 3 on brood mortality and egg hatch in *Solenopsis invicta*. J Invertebr Pathol 203: 108056.

Valles SM. 2023. Solenopsis invicta virus 3 infection alters foraging behavior in its host. Virology 581: 81–88.

Valles SM, Oi DH, Weeks RD, Addesso KM, Oliver JB. 2022. Field evaluation of Solenopsis invicta virus 3 against its host *Solenopsis invicta*. J Invertebr Pathol 191: 107767.

Valles SM, Shoemaker D, Wurm Y, Strong CA, Varone L, Becnel JJ, Shirk PD. 2013. Discovery and molecular characterization of an ambisense densovirus from South American populations of *Solenopsis invicta*. Biological Control 67: 431–439.

van Kempen M, Kim SS, Tumescheit C, Mirdita M, Lee J, Gilchrist CLM, Söding J, Steinegger M. 2024. Fast and accurate protein structure search with Foldseek. Nat Biotechnol 42: 243–246.

Van Oers MM, Herniou EA, Jehle JA, Krell PJ, Abd-Alla AMM, Ribeiro BM, Theilmann DA, Hu Z, Harrison RL. 2023. Developments in the classification and nomenclature of arthropod-infecting large DNA viruses that contain pif genes. Arch Virol 168: 182.

Varaldi J, Fouillet P, Ravallec M, López-Ferber M, Boulétreau M, Fleury F. 2003. Infectious behavior in a parasitoid. Science 302: 1930.

Varaldi J, Ravallec M, Labrosse C, Lopez-Ferber M, Boulétreau M, Fleury F. 2006. Artifical transfer and morphological description of virus particles associated with superparasitism behaviour in a parasitoid wasp. J Insect Physiol 52: 1202–1212.

Walker PJ, Siddell SG, Lefkowitz EJ, Mushegian AR, Adriaenssens EM, Alfenas-Zerbini P, Davison AJ, Dempsey DM, Dutilh BE, García ML, et al. 2021. Changes to virus taxonomy and to the International Code of Virus Classification and Nomenclature ratified by the International Committee on Taxonomy of Viruses (2021). Arch Virol 166: 2633–2648.

Wang J, Chitsaz F, Derbyshire MK, Gonzales NR, Gwadz M, Lu S, Marchler GH, Song JS, Thanki N, Yamashita RA, et al. 2023. The conserved domain database in 2023. Nucleic Acids Res 51: D384–D388.

Wang Y, Kleespies RG, Huger AM, Jehle JA. 2007. The Genome of Gryllus bimaculatus Nudivirus Indicates an Ancient Diversification of Baculovirus-Related Nonoccluded Nudiviruses of Insects. J Virol 81: 5395–5406.

Whitfield JB. 2002. Estimating the age of the polydnavirus/braconid wasp symbiosis. Proc Natl Acad Sci U S A 99: 7508–7513.

Whitfield ZJ, Dolan PT, Kunitomi M, Tassetto M, Seetin MG, Oh S, Heiner C, Paxinos E, Andino R. 2017. The Diversity, Structure, and Function of Heritable Adaptive Immunity Sequences in the Aedes aegypti Genome. Curr Biol 27: 3511–3519.e7.

Williams T, Bergoin M, van Oers MM. 2017. Diversity of large DNA viruses of invertebrates. J Invertebr Pathol 147: 4–22.

Wu X, Guarino LA. 2003. Autographa californica Nucleopolyhedrovirus orf69 Encodes an RNA Cap (Nucleoside-2′-O)-Methyltransferase. J Virol 77: 3430–3440.

Yang Z. 2007. PAML 4: Phylogenetic Analysis by Maximum Likelihood. Molecular Biology and Evolution 24: 1586–1591.

Zhang L, Ma R, Xu W, Billen J, He H. 2023. Comparative morphology and ultrastructure of the labial gland among castes of Camponotus japonicus (Hymenoptera: Formicidae). Arthropod Structure & Development 72: 101236.

Zhang Y-Z, Shi M, Holmes EC. 2018. Using Metagenomics to Characterize an Expanding Virosphere. Cell 172: 1168–1172.

Zhu S, Wang W, Wang Y, Yuan M, Yang K. 2013. The Baculovirus Core Gene ac83 Is Required for Nucleocapsid Assembly and Per Os Infectivity of Autographa californica Nucleopolyhedrovirus. J Virol 87: 10573–10586.

Anon. 2008. AcMNPV ac143 (odv-e18) is essential for mediating budded virus production and is the 30th baculovirus core gene. Virology 375: 277–291.

